# Daily repetitive magnetic stimulation induces microglia-dependent homeostatic synaptic remodeling

**DOI:** 10.64898/2026.05.24.727479

**Authors:** Paolo d’Errico, Christos Galanis, Dimitrios Kleidonas, Claudio Elgueta, Marlene Bartos, Andreas Vlachos

**Affiliations:** Department of Neuroanatomy, Institute of Anatomy and Cell Biology, Faculty of Medicine, University of Freiburg, Freiburg, Germany; Center BrainLinks-BrainTools, University of Freiburg, Freiburg, Germany; Center for Basics in Neuromodulation (NeuroModulBasics), Faculty of Medicine. University of Freiburg, Freiburg, Germany; Institute for Physiology I, Faculty of Medicine, University of Freiburg, Freiburg, Germany

**Keywords:** repetitive transcranial magnetic stimulation, microglia, synaptic remodeling, homeostatic plasticity, organotypic slice cultures

## Abstract

Therapeutic repetitive transcranial magnetic stimulation (rTMS) is delivered across repeated sessions to induce durable network-level adaptations, yet the cellular mechanisms governing these cumulative effects remain poorly understood. Using mouse organotypic brain cultures, we show that repeated daily intermittent theta burst stimulation with 900 pulses (iTBS900; 10 stimulation sessions over 2 weeks) induces a microglia-dependent adaptive synaptic response in pyramidal neurons. This response is characterized by reduced AMPAR-mediated miniature excitatory postsynaptic current (mEPSC) frequency and decreased dendritic spine density, consistent with adaptive remodeling of excitatory synaptic connectivity. Microglial depletion abolished this homeostatic response and instead resulted in increased synaptic strength and elevated spine density following repeated stimulation. Repeated daily iTBS900 also enhanced microglial uptake and degradative capacity while altering baseline and injury-induced microglial motility, indicating stimulation-dependent changes in microglial surveillance and effector functions. Together, these findings identify microglia as critical mediators of the homeostatic structural and functional adaptations induced by multi-session rTMS.

## INTRODUCTION

Repetitive transcranial magnetic stimulation (rTMS) is administered across repeated stimulation sessions to induce therapeutic network adaptations in neurological and psychiatric disorders (Lefaucheur et al., 2020). Mechanistic models of rTMS have historically focused on neurons, based on the assumption that magnetic stimulation primarily drives neuronal depolarization and synaptic plasticity. Consistent with this view, substantial *in vivo* and *in vitro* evidence demonstrates that even a single rTMS session can induce long-lasting structural and functional forms of plasticity (Galanis et al., 2025; Lu et al., 2025; Zhong et al., 2021). However, therapeutic rTMS is delivered across repeated stimulation sessions, and the cellular mechanisms that govern the cumulative network adaptations induced by such multi-session protocols are far from understood.

Emerging work identifies microglia as active contributors to experience- and activity-dependent synaptic remodeling including forms of plasticity linked to learning and memory (Badimon et al., 2020; Favuzzi et al., 2021; Nebeling et al., 2023; Weinhard et al., 2018). Beyond their classical role as innate immune sentinels, microglia continuously survey the brain parenchyma, respond dynamically to changes in neuronal activity, remodel synaptic structures, and release signaling molecules that shape circuit function (Coull et al., 2005; Kleidonas et al., 2023; Paolicelli et al., 2011; Wang et al., 2026). Recent work further suggests that microglia contribute to rTMS-induced plasticity and can respond to magnetic stimulation by modulating cytokine release and microglia–neuron communication (d’Errico et al., 2025; Eichler et al., 2023). Nevertheless, it remains unresolved whether microglia primarily support, constrain, or actively mediate the adaptive circuit remodeling induced by repeated therapeutic stimulation. In particular, the effects of multi-session rTMS on microglial surveillance behavior, motility, and phagocytic function remain poorly defined.

To address these questions, we subjected mouse entorhino-hippocampal slice cultures to repeated daily intermittent theta burst stimulation consisting of 900 pulses (iTBS900; 10 stimulation sessions over 2 weeks). Combining electrophysiology, dendritic spine analysis, live imaging, and pharmacological microglia depletion, we investigated how repeated iTBS900 shapes excitatory synapses and microglia function. We show that multi-session iTBS900 induces a microglia-dependent adaptive remodeling of excitatory synapses, characterized by reduced α-amino-3-hydroxy-5-methyl-4-isoxazolepropionic acid receptor (AMPAR)-mediated synaptic input and decreases dendritic spine density. In contrast, microglia depletion unmasks an opposite phenotype marked by enhanced synaptic strength and increased spine density. Moreover, repeated iTBS900 enhances microglial engulfment activity while altering baseline and lesion-induced motility. Together, these findings identify microglia as critical mediators of the homeostatic structural and functional adaptations induced by multi-session rTMS.

## RESULTS

### Repeated daily iTBS900 induces homeostatic reduction in excitatory synaptic connectivity

Three-week-old organotypic entorhino-hippocampal slice cultures (**Fig. 1A**) were stimulated using an iTBS protocol consisting of 900 pulses delivered at 65% maximum stimulator output (iTBS900; **Fig. 1B, C**). One stimulation session was applied daily for ten sessions over two weeks, with a weekend interval between stimulation blocks (**Fig. 1D**). Stimulation was delivered using a standard 70mm figure-of-eight coil positioned 7 mm beneath interface-configured slice cultures in a 35-mm Petri dish with 1 ml culture medium below the membrane insert. Cultures were returned to the incubator immediately after stimulation. Finite element modelling estimated a mean induced electric field strength of 20.6 V/m within the tissue (Fig. 1A; c.f. Galanis et al., 2024).

**Figure 1.**
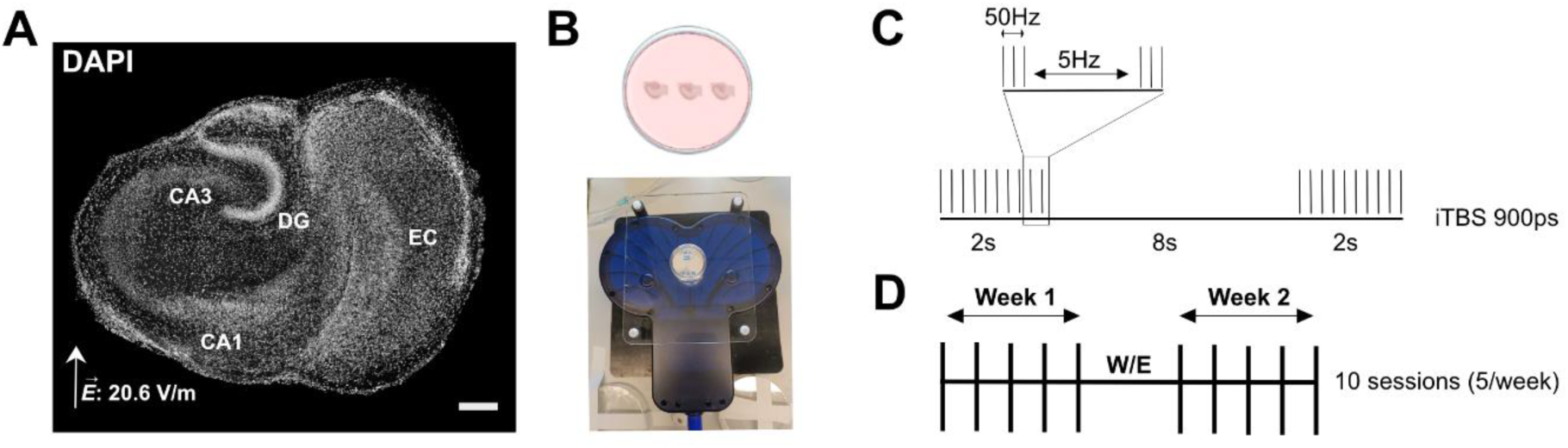
Experimental preparation and repeated daily iTBS stimulation paradigm. (**A**) Representative DAPI-counterstained slice culture and the corresponding computationally estimated electric field distribution during stimulation. DG, dentate gyrus; EC, entorhinal cortex; CA1 and CA3, Cornu Ammonis areas 1 and 3. Scale bar, 200 µm. (**B**) Schematic illustration and representative photograph of interface-configured entorhino-hippocampal slice cultures positioned above a 70-mm figure-of-eight coil for stimulation at 65% maximum stimulator output. (**C**) Intermittent theta burst stimulation protocol consisting of 900 pulses (iTBS900). (**D**) Multi-session stimulation schedule with one iTBS900 session per day over two weeks (5 sessions/week). W/E, weekend. Fig in B generated with Biorender.com.

Whole-cell recordings from CA1 pyramidal neurons (**Fig. 2A)** performed 2–4 h after the final stimulation session revealed no significant changes in AMPAR-mediated mEPSC amplitudes compared with age- and time-matched sham control. In contrast, mean mEPSC frequency was significantly reduced following repeated daily iTBS900 (**Fig. 2B**; Sham: 3.7 ± 0.35 Hz; rMS: 2.76 ± 0.30 Hz; Mann-Whitney U = 134, p < 0.05), indicating reduced excitatory synaptic input onto CA1 pyramidal neurons.

**Figure 2.**
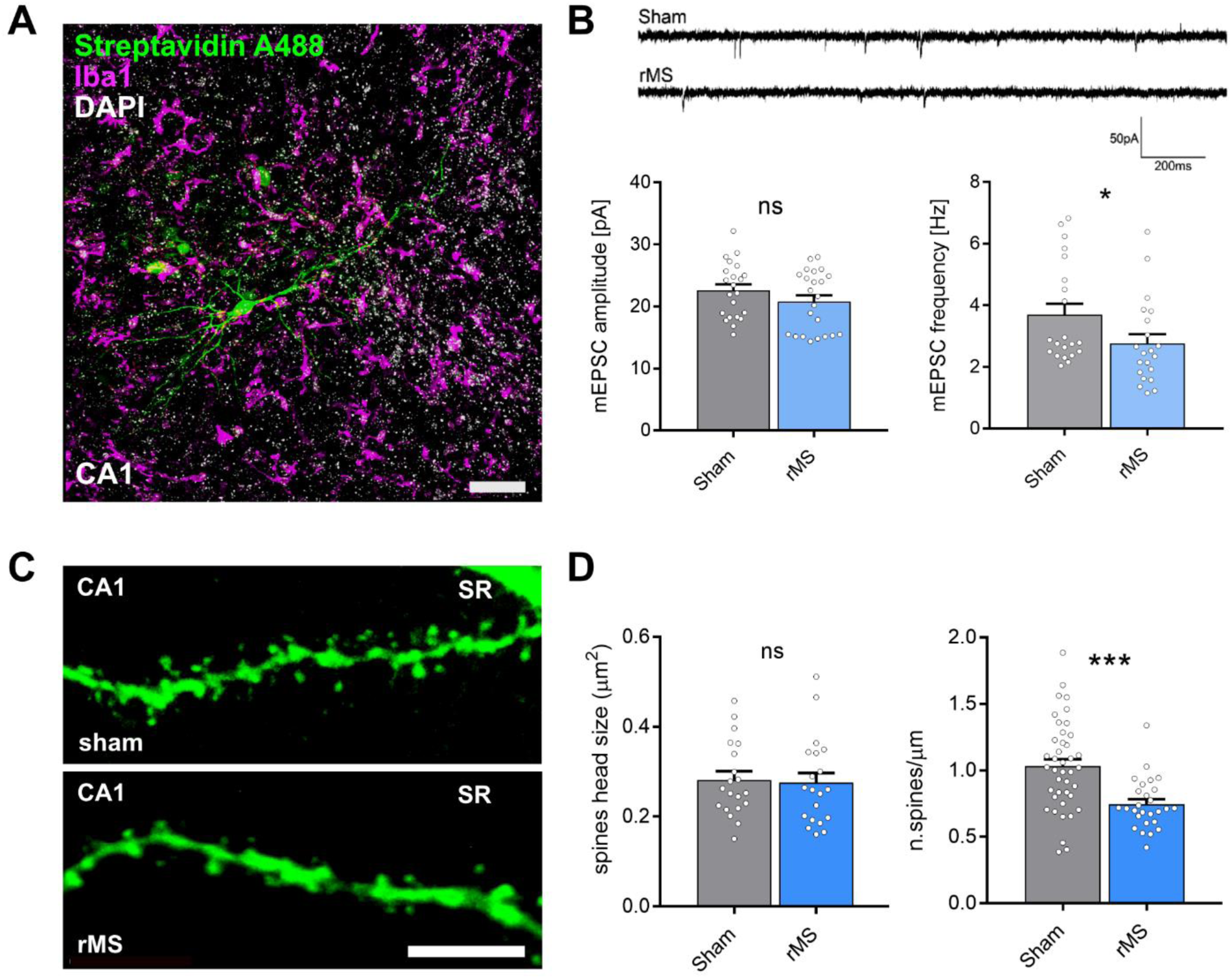
Repeated daily iTBS900 reduces miniature excitatory synaptic input and dendritic spine density in CA1 pyramidal neurons. (**A**) Representative confocal image of a biocytin-filled CA1 pyramidal neuron (Streptavidin-A488, green) and microglia (Iba1, magenta). Scale bar, 30 µm. (**B**) Representative traces (top) and quantification (bottom) of AMPAR-mediated miniature excitatory postsynaptic currents (mEPSCs) recorded from sham- and iTBS900-treated cultures (sham = 21 neurons from 8 cultures; iTBS900 = 21 neurons from 7 cultures; Mann-Whitney U = 134). Data represent mean ± SEM; each point represents an individual recorded neuron. (**C**) High-resolution confocal images of apical dendrites in SR from sham- and iTBS900-treated cultures. Scale bar, 5 µm. (**D**) Quantification of spine head size and spine density in SR dendritic segments (spine head size: sham = 20 dendritic segments from 6 cultures, iTBS900 = 20 dendritic segments from 5 cultures; spine density: sham = 41 dendritic segments from 6 cultures, iTBS900 = 27 dendritic segments from 5 cultures; Mann-Whitney U = 253). Data represent mean ± SEM, *p < 0.05, ***p < 0.001, ns: not significant.

To determine whether these functional changes were associated with structural alterations in excitatory synaptic connectivity, dendritic spines were quantified in recorded and biocytin-filled CA1 pyramidal neurons. Apical dendrites in *stratum radiatum* (SR), and in *stratum lacunosum moleculare* (SLM), as well as basal dendrites in *stratum oriens* (SO), were analyzed (**Fig. 2C; Fig. S1**). Repeated daily iTBS900 significantly reduced spine density selectively in SR, while spine head size remained unchanged (**Fig. 2D**; Sham: 1.03 ± 0.05 spines/µm; rMS: 0.74 ± 0.04 spines/µm; Mann-Whitney U = 253, p < 0.001). No significant changes were detected in SLM or SO (**Fig S1C, F**). These findings indicate a compartment-specific reduction in excitatory synaptic connectivity following repeated daily iTBS900.

### Microglia partial depletion reverses the effects of repeated daily iTBS900 on excitatory synapses

To assess whether microglia mediate the synaptic adaptations induced by repeated daily iTBS900, cultures were treated with the colony-stimulating factor 1 receptor (CSF1R) antagonist PLX3397 (100 nM) from 10 days in vitro (DIV10) until completion of the stimulation protocol (**Fig. 3A**; Coleman et al., 2020; Elmore et al., 2014). PLX3397 treatment reduced microglial density by 68% compared with vehicle-treated controls (**Fig. 3B, C**; −PLX: 411.3 ± 38.33 cells/mm^2^; +PLX: 133.2 ± 22.43 cells/ mm^2^; Mann-Whitney U = 4, p < 0.001) without affecting overall tissue viability, as assessed by propidium iodide (PI) staining. Exposure to NMDA (50 µM, 24 h) served as a positive control and induced robust PI fluorescence (**Fig. 3D–E**; DMSO: 0.428 ± 0.202 PI+ particles; PLX: 1.273 ± 0.468 PI+ particles; NMDA: 91 ± 20.56 PI+ particles; Kruskal-Wallis test, DMSO vs PLX: ns; DMSO vs NMDA: p < 0.001; PLX vs NMDA: p < 0.001).

**Figure 3.**
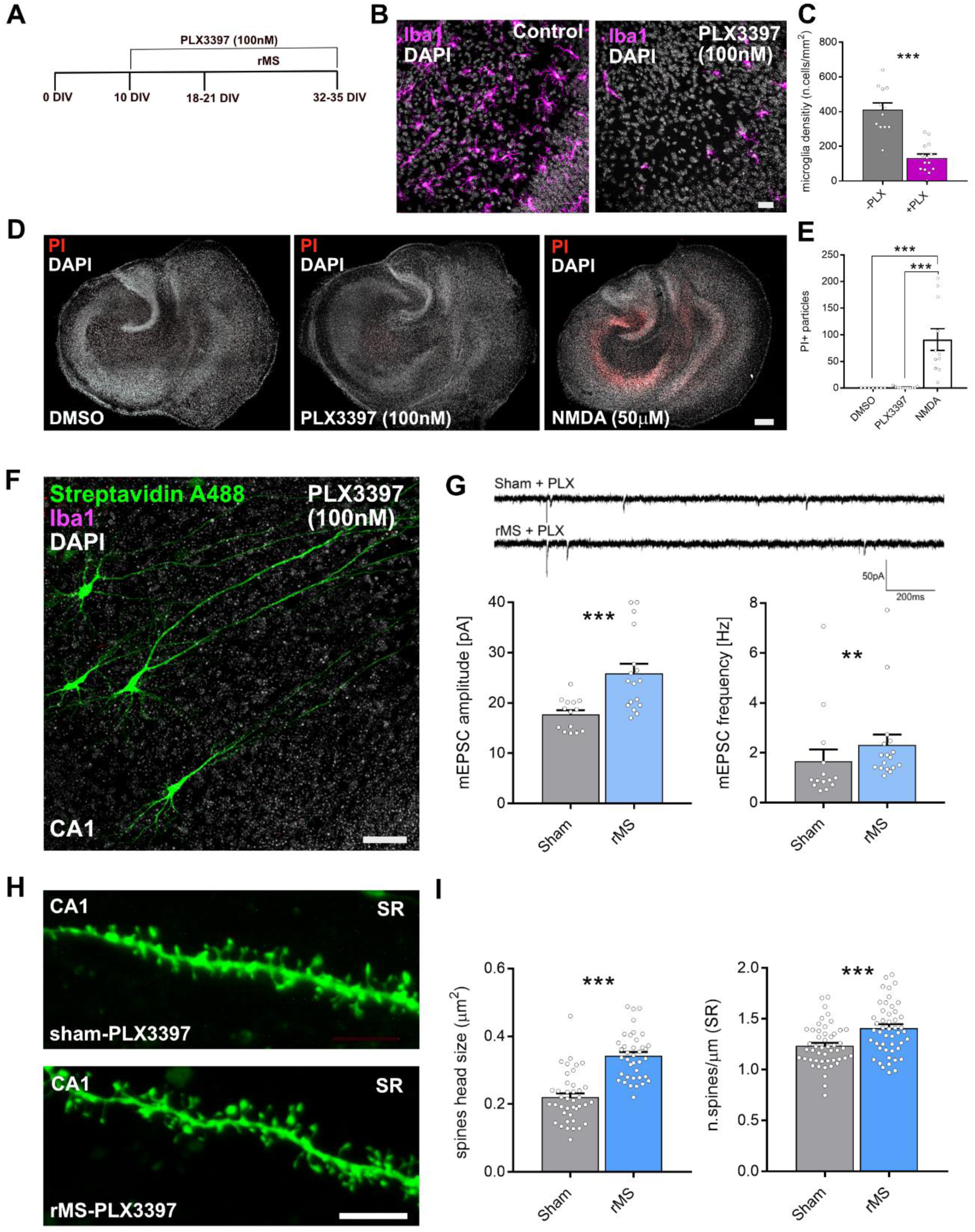
Microglia depletion alters iTBS900-induced functional and structural plasticity in CA1 pyramidal neurons. (**A**) Experimental timeline showing PLX3397 treatment (100 nM) and repeated daily ITBS900 in entorhino-hippocampal slice cultures. (**B**) Representative confocal images of microglia (Iba1, magenta) and nuclei (DAPI, white) in the CA1 region of vehicle-treated (-PLX; left) and PLX3397-treated (+PLX; right) cultures, illustrating effective microglia depletion. (**C**) Quantification of microglial cell density in vehicle- and PLX3397-treated cultures (-PLX = 12 cultures; +PLX = 14 cultures; Mann-Whitney U = 4; ***p < 0.001). **(D)** Representative confocal images of tissue cultures stained with propidium iodide (PI, red) and DAPI (white) under control conditions (left), following PLX3397 treatment (100 nM; middle), or after NMDA exposure (50 µM, 24 h; right). Scale bar, 200 µm. (**E**) Quantification of PI-positive particles in the indicated groups (DMSO = 7 cultures; PLX3397 = 11 cultures; NMDA = 11 cultures; Kruskal-Wallis test with Dunn’s multiple comparisons). (**F**) Representative confocal image of recorded and biocytin-filled CA1 pyramidal neurons (Streptavidin-A488, green) in a PLX3397-treated culture counterstained with DAPI-labeled nuclei (white). Scale bar, 30 µm. (**G**) Representative traces (top) and quantification (bottom) of AMPAR-mediated miniature excitatory postsynaptic currents (mEPSCs) recorded from sham- and iTBS900-treated PLX3397 cultures (sham = 14 neurons from 5 cultures; iTBS900 = 17 neurons from 6 cultures; Mann–Whitney U = 37; U = 53). (**H**) Representative images of CA1 apical dendrites in SR from sham-PLX3397 and rMS-PLX3397 conditions. Scale bar, 5 µm. (**I**) Quantification of spine head size and spine density in SR dendritic segments (spine head size: sham = 40 dendritic segments from 4 cultures, iTBS900 = 38 dendritic segments from 5 cultures; spine density: sham = 49 dendritic segments from 4 cultures, iTBS900 = 49 dendritic segments from 5 cultures; Mann–Whitney U = 148.5; U = 740). Data represent mean ± SEM, **p<0.01, ***p < 0.001.

In microglia-depleted cultures, repeated daily iTBS900 induced a markedly different synaptic response. Both AMPAR-mediated mEPSC amplitudes and frequencies were significantly increased following stimulation (**Fig. 3G**; sham amplitude: 17.72 ± 0.84 pA, iTBS900 amplitude: 25.9 ± 1.91 pA; Mann-Whitney U = 37, p < 0.001; Sham frequency: 1.656 ± 0.48 Hz; iTBS900 frequency: 2.32 ± 0.416 Hz; Mann-Whitney U = 53, p < 0.01), indicating enhanced excitatory synaptic strength. Consistent with these electrophysiological findings, dendritic spine analysis revealed increased spine head size and spine density, particularly in SR (**Fig. 3H–I; Fig. S1A–F**; sham size: 0.22 ± 0.01 μm^2^, iTBS900: 0.342 ± 0.01 μm^2^; Mann-Whitney U = 148.5, p < 0.001; Sham density: 1.23 ± 0.03 spines/μm, iTBS900 density: 1.41 ± 0.04 spines/μm; Mann-Whitney U = 740, p < 0.001). Thus, in the absence of microglia, repeated daily iTBS900 promoted structural and functional strengthening of excitatory synapses, consistent with previous observations following single rMS sessions (Lu et al., 2025; Vlachos, 2012).

To further assess structural remodeling, dendritic filopodia were quantified in SR dendrites (**Fig. S1H**). Filopodia were defined as thin elongated protrusions lacking spine heads (**Fig. S1G)** and are commonly considered immature spine-like structures. Repeated daily iTBS900 did not significantly alter filopodia density in either condition. However, PLX3397 treatment increased baseline filopodia density in SR dendrites (**Fig. S1H**; sham control: 0.048 ± 0.007 filopodia/μm; sham +PLX3397: 0.08 ± 0.009 filopodia/μm; Two-way ANOVA, p < 0.05), suggesting altered structural remodeling under microglia-depleted conditions.

Together, these findings demonstrate that microglia are required for the reduction in excitatory synaptic connectivity induced by repeated daily iTBS900. In the absence of microglia, the same stimulation paradigm instead promotes structural and functional strengthening of excitatory synapses.

### Repeated daily iTBS900 enhances microglial uptake and lysosomal processing

Microglia help refine neural circuits by selectively identifying and removing dendritic spines that are either overactive or functionally weak, thereby maintaining synaptic balance and efficiency (Cornell et al., 2022; Rueda-Carrasco et al., 2023). The opposing effects of repeated daily iTBS900 in microglia-intact and microglia-depleted cultures suggest an active role for microglia in the remodeling of excitatory synaptic connectivity following repeated stimulation. To assess microglial phagocytic activity, slice cultures were incubated with pHrodo™-labeled zymosan rhodamine particles, which light up upon acidification within lysosomal compartments. To evaluate uptake and subsequent processing dynamics, cultures were fixed 1 h (uptake) or 4 h (lysosomal processing) after particle exposure, and area of fluorescence signal within individual microglial soma was quantified (**Fig. 4A**). Microglia from iTBS900-stimulated cultures exhibited significantly greater particle-associated fluorescence after 1 h compared with sham controls (**Fig. 4B–C**; sham-1h: 0.63 ± 0.1 μm^2^; iTBS900-1h: 1.4 ± 0.2 μm^2^; Mixed Effects Model, p < 0.001). In sham cultures, area of fluorescence signal increased, although not significantly, from 1 h to 4 h, consistent with continued particle engulfment over time. In contrast, fluorescence area decreased between 1 h and 4 h in iTBS900-treated cultures (**Fig. 4B–C**; iTBS900-1h: 1.4 ± 0.2 μm^2^; iTBS900-4h: 0.7 ± 0.1 μm^2^; Mixed Effects Model, p < 0.05), suggesting enhanced lysosomal processing following repeated stimulation. Similarly, microglial soma area increased at 1 h after iTBS900 (**Fig. 4C**; sham-1h: 112.9 ± 3.89 μm^2^; iTBS900-1h: 138.2 ± 4.33 μm^2^; Mixed Effects Model, p < 0.01), but decreased between 1 h and 4 h in stimulated cultures (**Fig. 4C**; iTBS900-1h: 138.2 ± 4.33 μm^2^; iTBS900-4h: 105.6 ± 3.79 μm^2^; Mixed Effects Model, p < 0.01), indicating dynamic morphological changes associated with particle uptake and processing.

**Figure 4.**
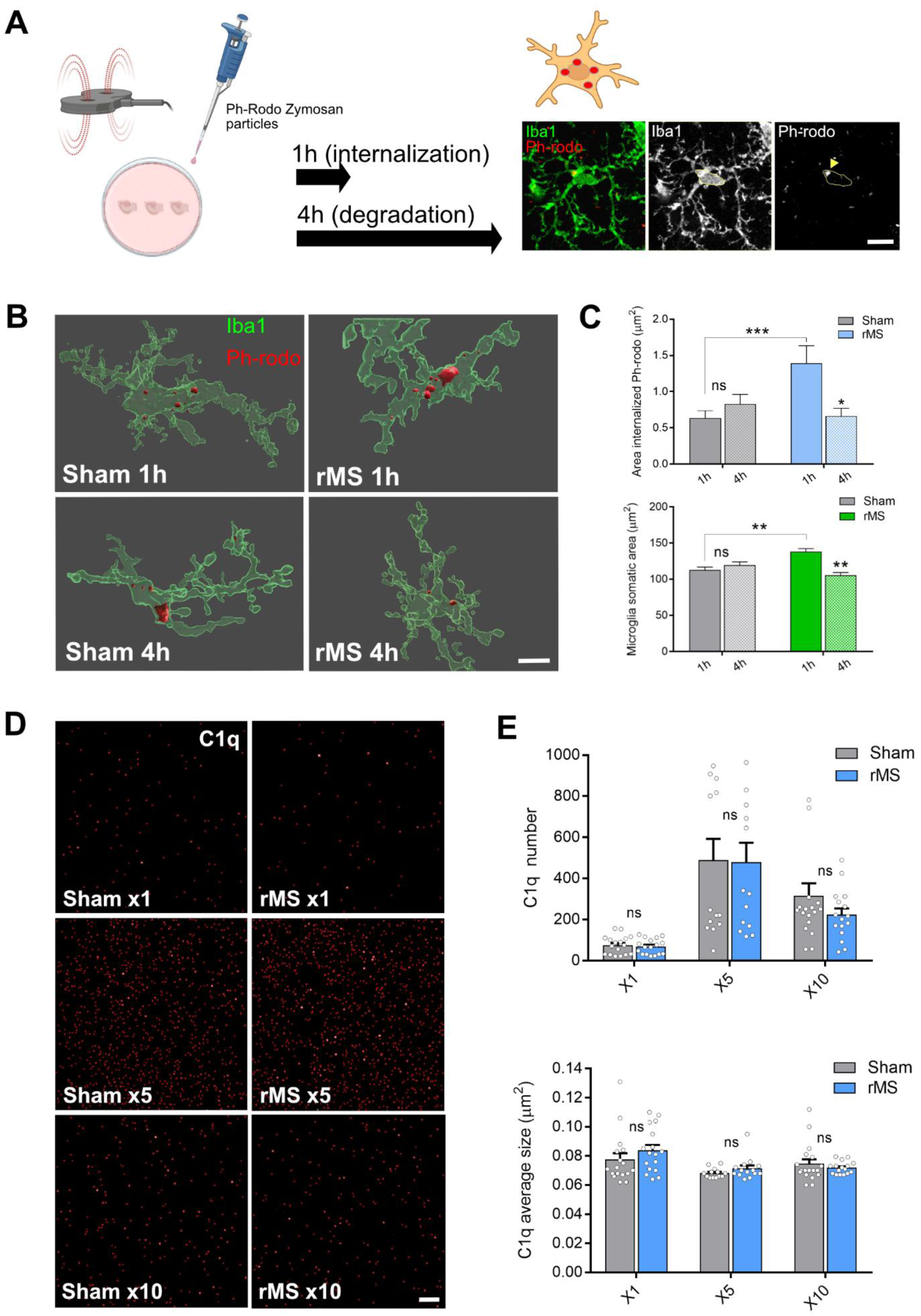
Repeated daily iTBS enhances microglial uptake and lysosomal processing. **(A)** Schematic illustration of the phagocytosis assay. Slice cultures were incubated with pHrodo™ Red Zymosan particles and fixed after 1 h to assess particle uptake or after 4 h to evaluate subsequent processing dynamics. Representative confocal images show Iba1-positive microglia (green) containing pHrodo-positive cargo (red). Scale bar, 5 µm. **(B)** Representative Imaris 3D reconstructions from confocal image stacks illustrating pHrodo™ Zymosan uptake in sham- and iTBS900-treated cultures at 1 h (top) and 4 h (bottom) after particle exposure. Scale bar, 10 µm. **(C)** Quantification of microglial pHrodo™ fluorescence intensity (top) and somatic area (bottom) at 1 h and 4 h in sham- and iTBS900-treated cultures (sham = 83–126 microglia from 9–14 cultures; iTBS900 = 104–136 microglia from 11–16 cultures; Mixed-effects model). **(D)** Representative confocal images of C1q puncta in SR following 1, 5, or 10 iTBS900 sessions. Scale bar, 5 µm. **(E)** Quantification of C1q puncta number (top) and mean puncta size (bottom) in sham- and iTBS900-treated cultures (sham = 14–19 cultures; iTBS900 = 15–17 cultures; Two-way ANOVA followed by Tukey’s multiple comparisons test). Data represent mean ± SEM, *p < 0.05, **p < 0.01, ***p < 0.001, ns: not significant.

The complement system represents a major pathway involved in microglia-mediated synaptic elimination, with C1q acting as an initiating recognition molecule (Hong et al., 2016; Stevens et al., 2007). To assess potential changes in complement-associated signaling during repeated stimulation, slice cultures were fixed after 1, 5, or 10 days of repeated daily iTBS900, and immunolabeled C1q puncta number and size were quantified in SR by immunofluorescence (**Fig. 4D**). C1q expression varied across time in culture, with highest levels observed during the fourth week *in vitro* in both sham- and iTBS900-treated cultures. However, repeated daily iTBS900 did not significantly alter either the number or size of C1q puncta at any analyzed time point (**Fig. 4E**). These findings suggest that repeated daily iTBS900 does not induce major changes in C1q-associated complement signaling under the present experimental conditions.

In summary, repeated daily iTBS900 stimulates microglia to increase particle uptake and accelerate lysosomal processing, accompanied by dynamic morphological changes, without significantly altering C1q-mediated complement signaling.

### Repeated daily iTBS900 differentially affects baseline and lesion-induced microglial motility

Microglia continuously survey the brain parenchyma through highly dynamic process motility and rapidly respond to local tissue injury (Davalos et al., 2005; Nimmerjahn et al., 2005). To assess whether repeated daily iTBS900 alters microglial surveillance dynamics, live imaging experiments were performed in slice cultures derived from Cx3cr1-GFP reporter mice following completion of the stimulation paradigm. Time-lapse recordings were acquired using confocal microscopy at 3-min intervals over 36 min under baseline conditions or using a two-photon microscope at 1-min intervals over 15 min following focal laser-induced injury. Process extension and retraction were quantified to determine movement velocity and total distance traveled (**Fig. 5A**).

**Figure 5.**
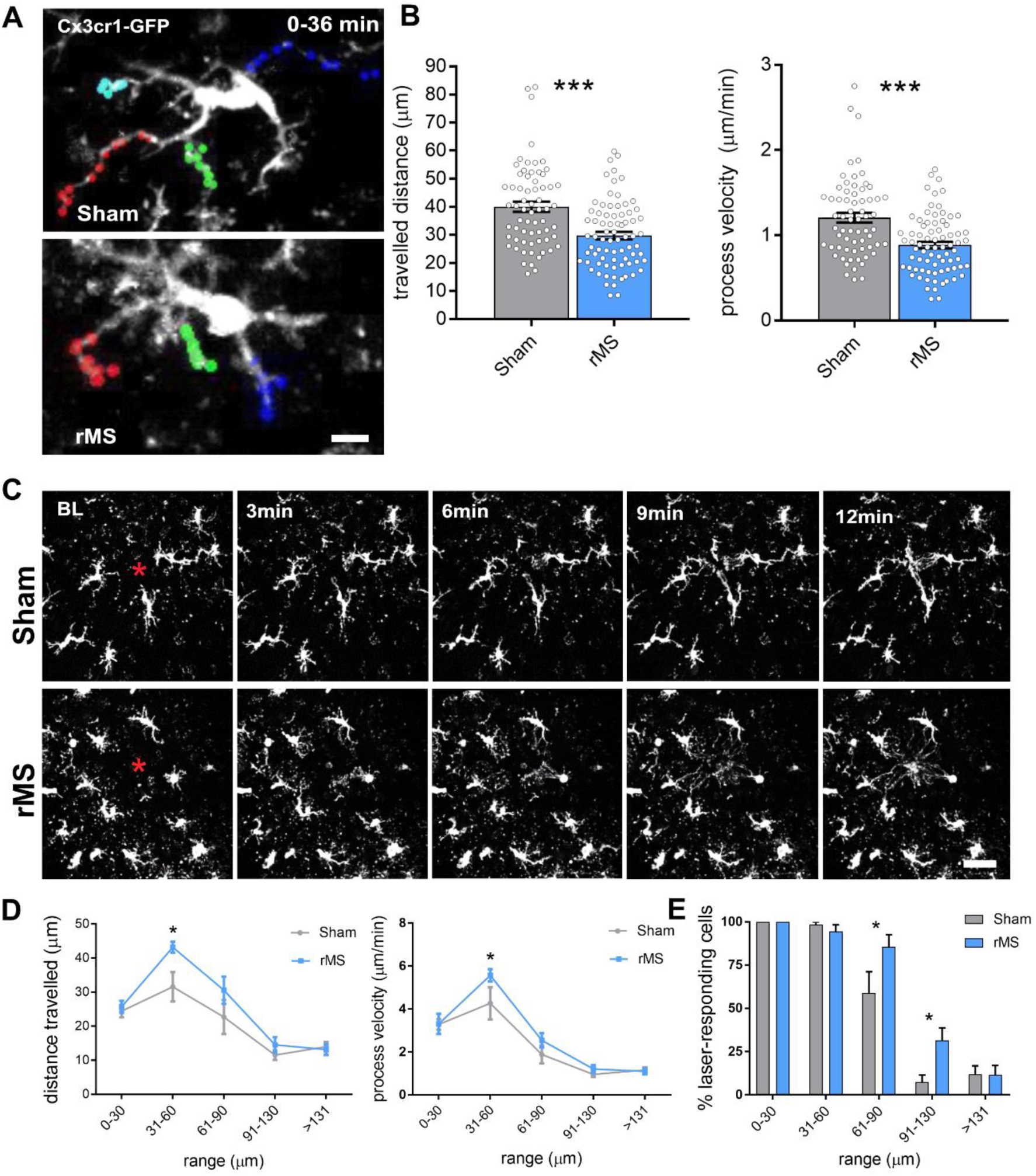
Repeated daily iTBS900 alters baseline and lesion-evoked microglial process motility. **(A)** Representative time-lapse images illustrating baseline microglial process motility in sham- and iTBS900-treated cultures. Colored points indicate tracked movements of individual microglial processes acquired at 3-min intervals over a total imaging period of 36 min. Scale bar, 20 µm. **(B)** Quantification of total traveled distance (left) and process movement velocity (right) under baseline conditions in sham- and iTBS900-treated cultures. Each point represents an individual tracked process (sham = 65 processes from 8 cultures; iTBS900 = 76 processes from 9 cultures; Mann–Whitney U = 1458; U = 1439). **(C–E)** *Ex vivo* two-photon imaging analysis of lesion-directed microglial process motility during the first 12 min following focal laser-induced injury (red asterisk) in sham- and iTBS900-treated cultures. BL, before lesion. Scale bar, 100 µm. **(D)** Quantification of total traveled distance (left) and process velocity (right) of microglial processes following laser-induced injury. (**E**) Percentage of laser-lesion responsive microglia as a function of distance from the lesion site (in D, E: sham = 13 lesions from 10 cultures; iTBS900 = 21 lesions from 14 cultures; Two-way ANOVA followed by Sidak’s multiple comparisons test). Data represent mean ± SEM, *p < 0.05, ***p < 0.001.

Following repeated daily iTBS900, microglial processes exhibited reduced baseline motility, reflected by decreases in both movement distance and velocity compared with sham-controls (**Fig. 5B**; sham distance: 40 ± 1.8 μm, iTBS900 distance: 29.8 ± 1.4 μm; Mann-Whitney U=1458, p < 0.001; Sham velocity: 1.2 ± 0.1 μm/min, iTBS900 velocity: 0.9 ± 0.04 μm/min; Mann-Whitney U=1439, p < 0.001). In contrast, focal laser-induced injury in SR using *ex vivo* two-photon imaging (**Fig. 5C**), revealed enhanced lesion-directed microglial motility following repeated daily iTBS900. Microglial processes displayed increased movement distance and velocity toward the laser-lesion site, particularly within 31–60 µm of the lesion core (**Fig. 5D**; distance: sham_31–60_: 31.6 ± 4.3 µm, iTBS900_31–60_: 43.2 ± 1.6 µm; Two-way ANOVA followed by Sidak’s multiple comparisons test, p < 0.05; Velocity sham_31–60_: 1.9 ± 0.4 µm/min, iTBS900_31–60_: 2.5 ± 0.3 µm/min; Two-way ANOVA followed by Sidak’s multiple comparisons test, p < 0.05). Quantification of responding microglia within regions located 61–130 µm from the lesion further demonstrated a higher proportion of responsive cells in iTBS900-treated cultures compared with sham controls (**Fig. 5E**; sham_61–90_: 58.8 ± 12.3 %cells, iTBS900_61–90_: 85.6 ± 6.9 %cells; Two-way ANOVA followed by Sidak’s multiple comparisons test, p < 0.05; Sham_91–130_: 7.38 ± 4.1 %cells, iTBS900_91–130_: 31.4 ± 7.3 %cells; Two-way ANOVA followed by Sidak’s multiple comparisons test, p < 0.05). Together, these findings indicate that repeated daily iTBS900 differentially modulates baseline surveillance motility and lesion-directed microglial responses.

## DISCUSSION

In the present study, we show that repeated daily iTBS900 induces a microglia-dependent adaptive remodeling of excitatory synaptic connectivity in CA1 pyramidal neurons. Repeated stimulation over two weeks reduced AMPAR-mediated mEPSC frequency and decreased dendritic spine density in SR, while leaving spine head size unchanged under microglia-intact conditions. In contrast, microglia depletion abolished this adaptive response and instead unmasked increased synaptic strength together with elevated spine density and spine size following the same stimulation protocol. These findings identify microglia as critical regulators of the long-term synaptic adaptations induced by repeated rTMS. Moreover, the enhanced microglial uptake and lysosomal processing observed after repeated iTBS900 suggest that microglia actively contribute to the refinement of excitatory synaptic connectivity during prolonged stimulation.

Our previous work demonstrated that single sessions of 10 Hz rMS or iTBS induce NMDAR-dependent potentiation of excitatory synaptic transmission and structural plasticity in CA1 pyramidal neurons (Eichler et al., 2023; Galanis et al., 2025; Lu et al., 2025; Vlachos, 2012). The present findings extend these observations by showing that repeated daily stimulation recruits additional adaptive mechanisms that fundamentally alter the direction of synaptic remodeling. Rather than promoting further potentiation, repeated iTBS900 induced a progressive reduction in excitatory synaptic connectivity that depended on the presence of microglia. Importantly, the opposite phenotype observed in microglia-depleted cultures indicates that microglia do not merely permit plasticity but actively shape the long-term outcome of repeated stimulation. Together with previous evidence that rMS rapidly modulates microglial cytokine signaling and neuron–microglia communication (Eichler et al., 2023), our findings support the idea that microglia integrate repeated stimulation over time and engage adaptive remodeling programs that constrain excessive excitation and stabilize network activity.

Repeated daily iTBS900 also altered several aspects of microglial behavior. Microglia exhibited enhanced uptake and lysosomal processing of pHrodo-labeled zymosan particles, indicating stimulation-dependent changes in engulfment and degradative capacity. At the same time, repeated stimulation reduced baseline process motility while enhancing lesion-directed responses following focal laser injury. Together, these findings suggest that repeated iTBS900 does not globally impair microglial function, but rather shifts microglia toward a distinct functional state characterized by altered surveillance dynamics and enhanced responsiveness to local perturbations. Although the precise signaling pathways remain unresolved, several candidate mechanisms may contribute to this form of adaptive remodeling, including cytokine-dependent signaling, purinergic pathways, fractalkine signaling, and complement-associated pathways involved in synapse elimination. Notably, however, we did not detect major changes in C1q puncta density or size under the present experimental conditions, suggesting that large-scale activation of complement-associated signaling is not required for the observed remodeling response.

The reduction in dendritic spine density was restricted to *stratum radiatum*, indicating compartment-specific remodeling of excitatory synapses. This observation is consistent with previous studies demonstrating that rTMS-induced structural and functional plasticity is preferentially expressed at proximal apical dendrites of CA1 pyramidal neurons (Hananeia et al., 2025; Lenz et al., 2015). Together with recent evidence for cooperative pre- and postsynaptic mechanisms in rTMS-induced synaptic plasticity (Galanis et al., 2025), the present findings suggest that repeated iTBS900 preferentially remodels synapses within strongly recruited input domains. Future pathway-specific analyses will be required to determine how local neuronal activity patterns are integrated with microglia-dependent remodeling processes during repeated stimulation. It should also be noted that the higher concentration of PLX3397 used in the present study induced subtle baseline alterations in excitatory synaptic properties, which need to be considered when interpreting stimulation-induced effects. Nevertheless, the central finding that repeated daily iTBS900 fails to induce homeostatic synaptic adaptations in the absence of microglia remained robust across experimental conditions.

Repeated stimulation sessions are a defining feature of therapeutic rTMS protocols used in neuropsychiatric disorders. The present findings provide a cellular framework for understanding how cumulative stimulation may progressively engage microglia-dependent adaptive mechanisms that stabilize network activity over time. Such homeostatic remodeling may contribute to preventing excessive excitation while maintaining the capacity for further circuit adaptation. At the same time, our findings further support the emerging concept that the neuroimmune state critically influences the outcome of brain stimulation. Indeed, previous work demonstrated that inflammatory conditions impair the expression of rTMS-induced synaptic plasticity, whereas anti-inflammatory signaling can restore plasticity competence (Lenz et al., 2020). Together, these observations suggest that microglial state and neuroimmune context represent key determinants of stimulation-induced network remodeling and may contribute to inter-individual variability in therapeutic responses to rTMS.

In summary, repeated daily iTBS900 induces a microglia-dependent homeostatic remodeling of excitatory synaptic connectivity accompanied by distinct changes in microglial functional behavior. Our findings identify microglia as active regulators of the long-term network adaptations induced by repeated magnetic stimulation and establish a mechanistic link between cumulative stimulation, microglial state, and sustained synaptic remodeling. These results provide further insight into the cellular basis of multi-session rTMS and highlight microglia as potential modulators of therapeutic brain stimulation responses.

## MATERIALS AND METHODS

### Ethics statement

Mice were maintained in a 12 h light/dark cycle with food and water available *ad libitum*. Every effort was made to minimize distress and pain of animals. All experimental procedures were performed according to German animal welfare legislation and approved by the appropriate animal welfare committee and the animal welfare officer of the University of Freiburg under license number X-21/01B and X-22/12B.

### Preparation of tissue cultures

Organotypic entorhino-hippocampal tissue cultures 300 µm thick, were prepared at postnatal day 4-5 from *C57BL/6J* and *Cx3cr1-GFP/wt* (here reported as *Cx3cr1+/-*) mice of both sexes as previously described (Del Turco and Deller, 2007). The tissue cultures were transferred for cultivation onto porous (0.4 µm pore size, hydrophilic PTFE) cell culture inserts with 30 mm diameter (Merck/Millipore, Darmstadt, Germany, Cat# PICM0RG50). The culturing medium consisted of 50 % (v/v) minimum essential medium (MEM), 25 % (v/v) basal medium eagle (BME), 25 % (v/v) heat-inactivated normal horse serum (NHS), 2 mM GlutaMAX, 0.65 % (w/v) glucose, 25 mM HEPES buffer solution, 0.1 mg/ml streptomycin, 100 U/ml penicillin and 0.15 % (w/v) bicarbonate. The pH of the culturing medium was adjusted to 7.3 and tissue cultures were allowed to mature for 18-20 days at 35 °C in a humidified atmosphere with 5 % CO_2_ before starting the rMS administrations for further two weeks. The culturing medium was replaced three times a week.

### Electric field modelling

Finite element method was used to create a three-dimensional mesh model consisting of two compartments, representing the bath solution and tissue cultures. The physical dimensions of the mesh model were based on the physical parameters of the *in vitro* settings, with a coil-to-Petri dish distance of 7 mm and the coil positioned below the culture. Electrical conductivities of 1.654 S/m and 0.275 S/m were assigned to the bath solution and culture, respectively. The rate of change of the coil current was set to 1.4 A/ms at 1% MSO and scaled up to higher stimulation intensities. Simulations of macroscopic electric fields were performed using SimNIBS (3.2.6) and MATLAB (2023a). A validated 70 mm MagStim figure-of-eight coil was utilized in all simulations. The mean value of the absolute E-field was extracted from the volume compartment of the tissue culture.

### Repetitive magnetic stimulation (rMS)

A 70-mm figure-of-eight coil (D70 Air Film Coil, Magstim) connected to a Magstim Super Rapid2 Plus1 stimulator (Magstim) was positioned 7 mm below the culture dish. Cultures were stimulated using the FDA-approved intermittent theta-burst stimulation (iTBS) protocol, consisting of bursts of three pulses at 50 Hz delivered every 200 ms (5 Hz), for a total of 900 pulses (10-s inter-train interval) at 65% of maximum stimulator output (MSO). This stimulation intensity induces an electric field of 20.65 V/m, as estimated by finite-element modeling, as previously described (Galanis, et al., 2024). Brain slice cultures were stimulated once daily over a period of two weeks, with a pause during weekends. Slices were oriented such that the induced electric field was aligned with the apical–basal axis of CA1 pyramidal neurons. Time-matched cultures that were not stimulated but otherwise treated identically served as sham-stimulated controls. Cultures were processed for experimental analyses two to four hours after the final rMS session.

### Whole-Cell Patch-Clamp Recordings

Whole-cell voltage-clamp recordings of CA1 pyramidal neurons were conducted at 35°C using a Luigs & Neumann setup (Luigs & Neumann, Germany) equipped with an Olympus BX51WI microscope (Olympus, Japan), a MultiClamp 700B amplifier and Digidata 1550B digitizer (Axon Instruments, Molecular Devices, USA), as described previously (Galanis et al., 2025). The bath solution consisted of 126 mM NaCl, 2.5 mM KCl, 26 mM NaHCO3, 1.25 mM NaH2PO4, 2 mM CaCl2, 2 mM MgCl2, and 10 mM glucose and was saturated with 95% O2/5% CO2. 10 μM D-APV and 0.5 μM TTX were added in the bath solution to isolate miniature α-amino-3-hydroxy-5-methyl-4-isoxazolepropionic acid (AMPA) receptor-mediated excitatory postsynaptic currents (mEPSCs). Patch pipettes were pulled from borosilicate glass capillaries (1.5 mm outer diameter; Harvard Apparatus, USA) and filled with an internal solution containing 126 mM K-gluconate, 4 mM KCl, 4 mM ATP-Mg, 0.3 mM GTP-Na2, 10 mM phospho-creatine, 10 mM HEPES, and 0.1% (w/v) biocytin (pH 7.25 with KOH, 290 mOsm with sucrose). Patch pipettes had resistances of 3–5 MΩ for both current- and voltage-clamp recordings, and were matched across conditions. The neurons were recorded at −70 mV. Liquid junction potential was not corrected in these experiments. Signals were sampled at 10 kHz with a 6 kHz low-pass Bessel filter.

### Pharmacology for microglia depletion

To deplete microglia in tissue cultures, the colony-stimulating factor 1 receptor (CSF1R) inhibitor PLX3397 (100 nM; Axon MedChem, Groningen, The Netherlands; Cat#2501) dissolved in DMSO was added to the culture medium starting at DIV 10 and maintained throughout the entire cultivation period. Control cultures received an equivalent volume of DMSO.

### Microglia Phagocytic assay with pHrodo™ Red Zymosan particles

One hour after the final rMS session, pHrodo™ Red Zymosan BioParticles™ (200 nM; Invitrogen; Cat# P35364), diluted in 2 µL of incubation medium, were applied directly onto the slice cultures and incubated for either 1 or 4 h. Slices mounted on membrane inserts were then rapidly washed in 0.01 M PBS and fixed in 4% paraformaldehyde (PFA) for 1–2 h at room temperature. Following fixation, slices were transferred to PBS and stored at 4 °C until further processing.

### Immunostaining and imaging

Tissue cultures were fixed for 1-2 hours in a solution of 4% PFA in PBS at room temperature. After fixation, cultures were rinsed three times in PBS (0.01 M, pH 7.4) and consecutively incubated for 1 h with 10% NGS in 0.5% Triton X-100 containing PBS to reduce nonspecific staining and to increase antibody penetration. Subsequently, cultures were incubated overnight at 4°C with the primary antibodies diluted in PBS with 10% NGS and 0.1% Triton X-10. The primary antibodies used are the following: guinea pig anti-Iba1 (1:500; Synaptic System; Cat#HS-234308), mouse anti C1q (1:500; Abcam; Cat#ab-182451). Sections were washed and incubated overnight at 4°C with appropriate AlexaFluor dye-conjugated secondary antibodies (1:1000; Invitrogen) in PBS with 10% NGS, 0.1% Triton X-100. DAPI nuclear stain (1:3000 in PBS for 20 min; Fisher Scientific, Cat#62248) was used to visualize cytoarchitecture. Cultures were washed, transferred onto glass slides, and mounted for visualization with DAKO anti-fading mounting medium.

For *post hoc* visualization of recorded neurons, biocytin filled cells were labeled with the appropriate Alexa-conjugate streptavidin (Thermo Fisher Scientific; 1:1000; in 0.01 M PBS with 10% NGS, 0.1% Triton X-100) for 2 h and counterstained with DAPI. Confocal images at a resolution of 1024 x 1024 pixel were acquired using a Leica TCS SP8 laser scanning microscope equipped with 20x (NA 0.75), 40x (NA 1.30) and 63x (NA 1.40) oil-submersion objectives.

### Live-cell imaging of surveillance microglia process motility

To assess microglial process dynamics in steady state conditions after two-week rMS, live-cell imaging of tissue cultures from Cx3cr1+/- mice was performed at a Zeiss LSM800 confocal microscope. Filter membranes with 2-3 cultures were transferred in a in house 3D- printed dish containing circulating aCSF under continuous oxygenation (5% CO_2_/95% O_2_). The cultures were kept at 35°C during the whole imaging procedure. Two-to-four hours after the last rMS session, GFP+ microglia process motility was recorded in SR hippocampal area every 3 min for a total of 36 min at a depth of 30-35 µm from the slice surface with 2-µm *z* increments (total stack of 40 µm) and 512 x 512 pixels resolution using a 10x water immersion objective (NA 0.3) objective with 2x optical zoom. In each slice 1-2 sites were recorded.

### 2PM-laser lesion

Focal lesions were induced in organotypic cultures continuously perfused with ACSF containing (in mM): 124 NaCl, 2.5 KCl, 24 NaHCO₃, 1.2 NaH₂PO₄, 2 MgSO₄, 2 CaCl₂, and 12.5 glucose, warmed to ∼32 °C. The aCSF had an osmolarity of ∼310 mOsm and was continuously bubbled with 95% O₂ and 5% CO₂ to maintain a pH of 7.43. A galvanometric two-photon laser scanning system (Femto2D-Alba, Femtonics Ltd., equipped with a Chameleon Ultra II laser, Coherent) was used to induce focal lesions and to record time-lapse images of GFP-labeled microglia in the CA1 *stratum radiatum*. After acquiring a baseline z-stack centered at a depth of 30-35 µm from the slice surface (300 × 300 µm, 10 sections, 2 µm spacing), a spiral stimulation pattern with a 5 µm radius was applied to induce a lesion (15 s, 70 mW, 880 nm) at a location devoid of microglial soma or processes. The same z-stack was subsequently acquired every minute for a total of 15 min. Data were denoised and smoothed using a Gaussian filter (MES software, Femtonics Ltd.).

### Quantification of process microglia movement

Microglia process motility in steady state conditions was tracked manually using the Manual Tracking Plugin of ImageJ. At least three processes per cell were followed along the 36 min recording and the mean speed and distance traveled calculated for each microglial process. In laser lesion experiment, the extension of individual microglia process was tracked manually and measured using ImageJ considering the initial distance (at *t*=0 min) and the final distance (at *t*=15 min) from the center of the laser lesion. The mean speed and distance traveled were calculated for each microglial process. The percentage of responsive microglia was quantified by counting cells that showed at least one process moving toward the lesion site. The positions of the processes at *t*= 0 min were divided into four ranges: 0-30 µm, 31-60 µm, 61-90 µm and 91-120 µm (see the results above).

### Statistical analysis

GraphPad Prism 7 and R-studio was used for statistical analysis. Mann-Whitney U test, Two-way ANOVA followed by Sidak’s or Tukey’s multiple comparisons test, Kruskal-Wallis test with Dunn’s multiple comparisons test and Mixed-effects model were used to assess statistical differences between experimental groups.

## AUTHOR CONTRIBUTIONS

PD: conceptualization, investigation, methodology, data analysis and quantifications, data curation, visualization, writing original draft, manuscript review & editing. CG: electrophysiology recordings & analysis, manuscript review and editing. DK: electrophysiology recordings & analysis, manuscript review and editing. CE: laser lesion and 2PM imaging, manuscript review and editing. MB: manuscript review and editing. AV: conceptualization, data curation, project supervision, funding acquisition, project administration, writing original draft, manuscript review & editing.

## Acknowledgements

This work was supported by the Federal Ministry of Education and Research, Germany (BMBF, 01GQ2205A) and Deutsche Forschungsgemeinschaft (DFG, CRC/TRR 167-Project-ID 259373024 B14)

## ACKNOWLEDGEMENT

The authors thank Zsolt Turi for support with finite element modelling and Emina Deumic for her skilful assistance in tissue culturing.

---

**Supp. Figure.**
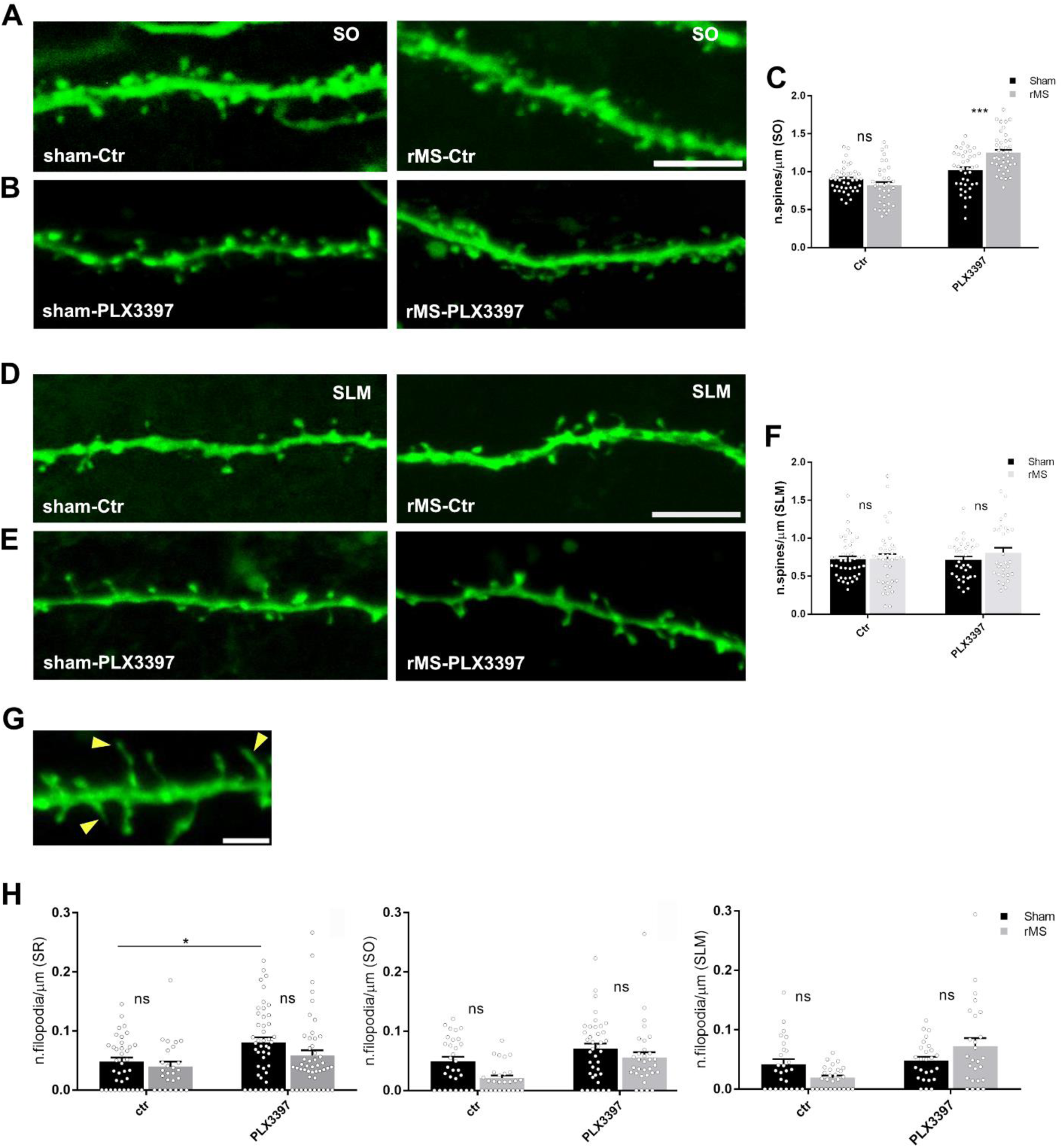
Repeated daily iTBS900 does not induce major structural changes in dendritic spine density of CA1 pyramidal neurons. (**A-B**) High-resolution confocal image of a biocytin-filled CA1 pyramidal neuron (Streptavidin-A488, green) in SO in control (A) and PLX3397-treated (B) conditions. Scale bar, 5 µm. (**C**) Graphs show spine density in SO in the respective groups (sham-Ctr = 43 dendritic segments from 5 cultures, iTBS900-Ctr =35 dendritic segments from 5 cultures; sham-PLX = 36 dendritic segments from 4 cultures, iTBS900-PLX = 41 dendritic segments from 5 cultures; Two-way ANOVA followed by Sidak’s multiple comparisons test). (**D-E**) High-resolution confocal image of a biocytin-filled CA1 pyramidal neuron (Streptavidin-A488, green) in SLM in control (D) and PLX3397-treated (E) conditions. Scale bar, 5 µm. (**F**) Graphs show spine density in the respective groups (sham-Ctr = 42 dendritic segments from 6 cultures, iTBS900-Ctr = 38 dendritic segments from 5 cultures; sham-PLX = 35 dendritic segments from 4 cultures, iTBS900-PLX = 29 dendritic segments from 5 cultures; Two-way ANOVA followed by Sidak’s multiple comparisons test). (**G**) High-resolution confocal image of a biocytin-filled CA1 pyramidal neuron (Streptavidin-A488, green) with representative filopodia (yellow head-arrows). Scale bar, 5 µm. (**H**) Graphs show filopodia density quantified in SR (left), SO (middle) and SLM (right) in the respective groups. (SR: sham-Ctr = 37 dendritic segments from 6 cultures, iTBS900-Ctr = 26 dendritic segments from 5 cultures; sham-PLX = 35 dendritic segments from 4 cultures, iTBS900-PLX = 29 dendritic segments from 5 cultures; SO: sham-Ctr = 30 dendritic segments from 6 cultures, iTBS900-Ctr = 27 dendritic segments from 4 cultures; sham-PLX = 38 dendritic segments from 4 cultures, iTBS900-PLX = 32 dendritic segments from 5 cultures; SLM: sham-Ctr = 32 dendritic segments from 6 cultures, iTBS900-Ctr= 31 dendritic segments from 4 cultures; sham-PLX = 26 dendritic segments from 4 cultures, iTBS900-PLX = 27 dendritic segments from 5 cultures; Two-way ANOVA followed by Sidak’s multiple comparisons test). Data represent mean ± SEM, *p<0.05, ***p<0.001, ns: not significant.

